# Auxin export from proximal fruits drives arrest in competent inflorescence meristems

**DOI:** 10.1101/542662

**Authors:** Alexander Ware, Catriona H. Walker, Jan Šimura, Karin Ljung, Anthony Bishopp, Zoe Wilson, Tom Bennett

**Author notes:** These authors contributed equally to this work (Alexander Ware, Catriona H. Walker).

## Abstract

A well-defined set of regulatory pathways control entry into the reproductive phase in flowering plants [1]. Conversely, little is known about the mechanisms that control the end of the reproductive phase (‘floral arrest’), despite this being a critical process for optimising fruit and seed production. Complete fruit removal or lack of fertile fruit-set in male sterile mutants, for example *male sterile1* (*ms1*), prevents timely floral arrest in the model plant Arabidopsis [2]. These observations formed the basis for Hensel and colleagues’ model in which end-of-flowering was proposed to result from a cumulative fruit/seed-derived signal that caused simultaneous ‘global proliferative arrest’ (GPA) in all inflorescences [2]. Recent studies have suggested that end-of-flowering involves gene expression changes at the floral meristem which are at least in part controlled by the *FRUITFULL-APETELA2* pathway [3,4], however there is limited understanding of how this process is controlled and the communication needed at the whole plant level. Here, we provide new information providing a framework for the fruit-to-meristem (F-M) communication implied by the GPA model [5]. We show that floral arrest in Arabidopsis is not ‘global’ and does not occur synchronously between branches, but rather that the arrest of each inflorescence is a local process, driven by auxin export from fruit proximal to the inflorescence meristem (IM). Furthermore, we show that inflorescence meristems are only competent for floral arrest once they reach a certain developmental age. Understanding the regulation of floral arrest is of major importance for the future manipulation of flowering to extend and maximise crop yields.

## RESULTS & DISCUSSION

To test the model proposed by Hensel et al, in which fruit number increases until reaching a cumulative threshold that causes inflorescence arrest, we performed a variety of fruit removal experiments in Arabidopsis. Firstly, we removed all open flowers on every branch continuously. This treatment prevented IM arrest on every branch, demonstrating that timely IM arrest indeed requires fertile fruit (**Fig. 1A,C**). Subsequently, we performed localised continuous flower removal on secondary cauline inflorescences only, and observed timely IM arrest (relative to control plants) everywhere on the plant apart from the treated inflorescences (**Fig. 1D, E**). These data show that floral arrest is not a ‘global’ systemic process, but rather consists of independent locally-regulated arrest of individual IMs.

**Figure 1.**
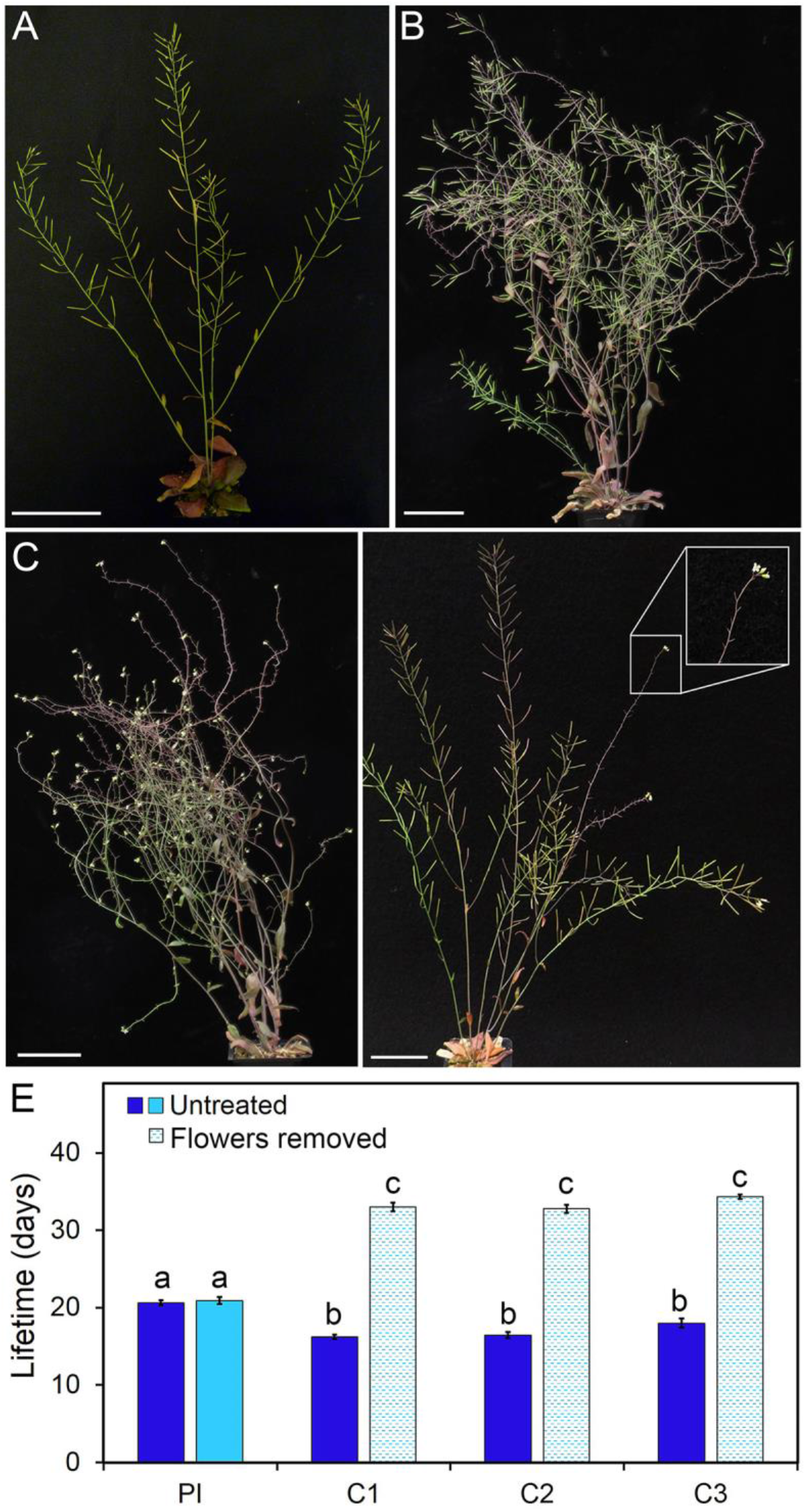
Inflorescence meristem arrest is a locally regulated process. (**A-D**) Inflorescence meristem arrest is delayed by continuous flower removal. Continuous daily removal of flowers across all inflorescences delays floral arrest in wild-type Arabidopsis (**A, C**), but when treatment is ended fruits develop, and arrest occurs within a few days (**B**). Local flower removal prevents arrest of individual inflorescences, but has no systemic effect (**D**). (**E**) Inflorescence lifetime in response to local flower removal. Open flowers were removed from secondary cauline inflorescences (C1, C2, C3) every 1-2 days until 17 days post bolting (DPB), whereupon open flowers were removed daily. Inflorescence lifetime was significantly extended in secondary inflorescences where flowers were removed (hatched light blue bars), relative to untreated plants (dark blue bars). However, the lifetime of untreated inflorescences, including the primary inflorescence (PI) on the same plants was not altered (light blue bar). *n* = 11-12, bars indicate s.e.m. Bars with the same letter are not statistically different from each other (ANOVA, Tukey HSD test).

A key tenet of Hensel et als’ GPA model is that floral arrest is synchronous. However, if IM arrest is locally-regulated, we questioned how it could also be synchronous across the plant. We therefore performed a detailed re-assessment of floral arrest in Arabidopsis. By tracking the lifetime of each inflorescence in a cohort of Col-0 plants, we found that IM arrest across plants is not synchronous, with on average ~5 days between arrest of the first and last IMs (**Fig 2B, Fig S1, Table S1**). The timing of arrest followed a clearly defined basipetal wave, with the primary inflorescence (PI) and the cauline (C) inflorescences arresting first, and the rosette (R) inflorescences arresting later (**Fig 2B**).

**Figure 2.**
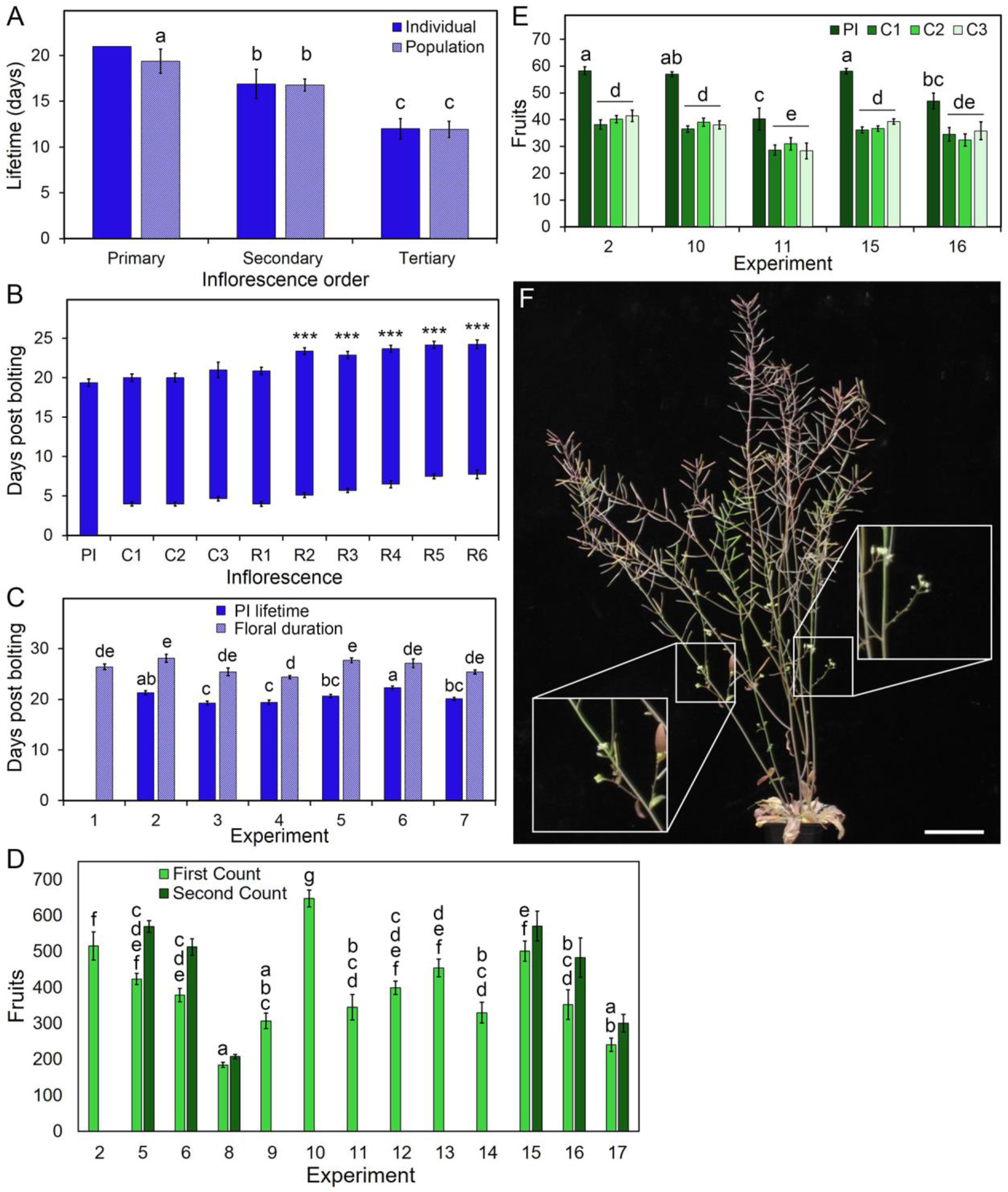
Inflorescence meristem arrest is a time-dependent process. (**A**) Timing of inflorescence meristem activation and arrest across different branches. For the primary (PI) and secondary inflorescences (including cauline (C1, C2 etc.) and rosette (R1, R2, etc.) inflorescences in a population of Col-0, the mean time after the floral transition, until IM activation and arrest, were measured. Each bar is the mean of 3-8 plants, depending on which plants had which inflorescence type. Any inflorescence type occurring on two or fewer plants was excluded from analysis. Error bars indicate s.e.m. Bars with the same letter are not statistically different from each other (ANOVA, Tukey HSD test). (**B**) Mean lifetime, from activation to arrest, of different classes of inflorescences, in a single Col-0 plant, and across a population of Col-0 plants. Bars indicate standard deviation. For the population, n=8 plants. Asterisks indicate statistically significantly different time of arrest from the primary inflorescence (ANOVA, Dunnett’s test, n=3-8, * =p<0.05, ** = p<0.01, *** = p<0.001). (**C**) Lifetime of the PI, and time from floral transition to *initial* floral arrest (floral duration), in Col-0 plants grown in long days (16h light/8h dark) in 6 experiments. *n*=10-12, bars indicate s.e.m. Bars with the same letter are not significantly different from each other (ANOVA, Tukey HSD test). (**D**) Mean total fruit production in long day-grown Col-0 plants across 13 separate experiments when initially counted (light green bars). In experiments 2 and 9-14, fruit numbers were only measured at the end of the experiment, and the ‘first count’ (light green bars) therefore includes any re-flowering that occurred. In experiments 5, 6, 8 and 15-17, we explicitly measured the number of fruit before re-flowering (light green bars) and again after re-flowering (‘second count’, dark green bars), n=8-18 depending on experiment, bars indicate s.e.m. Bars with the same letter are not significantly different from each other (ANOVA, Tukey HSD test). (**E**) Number of fruits on the primary inflorescence (‘PI’) and secondary cauline (C1, C2, C3) inflorescences in long day-grown across five experiments. *n*=6-13 depending on experiment, bars indicate s.e.m. Statistical tests relate to comparison of PI fruit, and comparison of combined C1, C2 and C3 fruit. Bars with the same letter are not significantly different from each other (ANOVA, T ukey HSD test). (**F**) Photograph showing re-flowering in Col-0, with new branches produced after initial floral arrest highlighted in white boxes. Scale bar = 5cm.

This corresponds to the similar basipetal wave of inflorescence activation observed earlier in the experiment (**Fig 2B, Fig S1**). We also observed that each class of inflorescence (e.g. primary, secondary, tertiary) had a distinctive lifetime, which was consistent both between inflorescences on the same plant, and between inflorescences on different plants (**Fig. 2A**). Indeed, we observed that, between different experiments run under similar conditions (**Table S1**), the lifetime of the PI in Col-0 was also consistently 20.5 (± 2) days) (**Fig. 2C**). We also observed that the timing of global floral arrest was also consistent between experiments, occurring within a window of 3 days either side of 25 days post-bolting (dpb) (**Fig. 2C**).

Conversely, we found that different numbers of fruit were produced between experiments run under similar conditions (**Table S1**), both in total (**Fig. 2D**), and on specific inflorescences (**Fig. 2E**). We found that the number of fruits produced on the PI was highly variable, ranging from ~30 to ~60 (**Fig 2E**), and that the ratio of fruits on the PI and secondary inflorescences varied between experiments (**Fig. 2E**). However, the number of fruits produced on the secondary cauline inflorescences was similar within experiments, indicating that the fruit production per inflorescence is not random, but tightly regulated (**Fig. 2E**). We also observed an additional phenomenon of ‘re-flowering’ in a number of experiments, whereby after arrest of most or all inflorescences, previously dormant axillary buds would re-activate, giving rise to new inflorescences (**Fig. 2F**). The re-initiation of flowering was not observed in all plants, nor indeed in all experiments, but it was a common occurrence. The number of additional fruits produced through re-flowering varied greatly between experiments, but was generally greatest in those experiments with a higher initial fruit production (**Fig. 2D**). It is thus likely that the vigour of re-flowering within an individual is controlled through the size of the plant at the time of the initial floral arrest. Larger plants with a greater number of inflorescences will typically have produced a greater number of dormant axillary buds, which can be re-activated during the re-flowering process.

Collectively, our data suggest that global cumulative fruit production is not the key factor driving floral arrest. Very different numbers of fruit are produced in different experiments before initial floral arrest, and more fruit can be produced as a result of re-flowering. Nor can we observe a correlation between the arrest of individual IMs and the number of fruit produced in our extensive datasets. Instead, our data suggest that time is a more relevant factor governing floral arrest, and more specifically the developmental time since the floral transition. Here, we define developmental age as the life-cycle stage of the inflorescence, as opposed to its absolute age, and recognise that the relationship between developmental and absolute age may vary due to environmental influences or differences in growth history. Our observations support a model in which each class of IM has a determined lifetime upon reaching a certain ‘developmental age’, become responsive to arrest-inducing signals. Thus, floral arrest thus occurs when all active IMs reach the end of their lifetime, and its timing is largely a reflection of the timing of IM activation. In instances where IM activation is synchronous, floral arrest may also be near-simultaneous. To reconcile the requirement of fruit to trigger IM arrest (**Fig. 1**) with the role of developmental age in regulating the timing of IM arrest, we hypothesised that at least some fertile fruit are required to allow temporally-competent IMs to undergo arrest.

To test this hypothesis we used a dexamethasone-inducible *MS1:MS1-GR* construct to restore fertility to the *ms1-1* mutant (L*er* background) 12 days post anthesis of the first flower (dpa). Restoring MS1 expression facilitated self-fertilisation and fertile fruit formation, and resulted in the timely arrest of inflorescences (**Fig. 3A**). However, the number of fertile fruit per inflorescence was still much lower than in wild-type plants (**Fig. 3A**). Similarly, if we removed flowers continuously from inflorescences beyond their normal lifetime, and then allowed plants to recover, each IM arrested within a few days, despite having produced only a small number of fruits (**Fig. 1B**). These data support a model in which fertile fruit-set has a role in regulating IM arrest, but do not support a model in which IM arrest is regulated by cumulative seed set, as in both cases meristems arrest with comparatively few fruit. To gain an increased understanding of the role of fruit in IM arrest, we performed partial and differential removal of flowers from inflorescences. ‘Early’ plants had all flowers removed, until the around 30 flowers had been produced by the PI, and were then allowed to continue flowering normally. ‘Late’ plants were allowed to flower as normal until around 30 flowers had opened on the PI; subsequently all open flowers were removed from the plant. Despite producing far fewer fruit than control plants (**Fig. 3B**), the PI of ‘early’ plants underwent arrest at the same time as untreated plants (**Fig. 3C**). Conversely, despite producing the same number of fruit as ‘early’ plants (**Fig. 3B**), ‘late’ plants did not undergo timely arrest (**Fig. 3C**). However, when flower removal treatment was ended in ‘late’ plants ~30 dpb, the IMs produced a small number of fruits and arrested. These data conclusively demonstrate that the role of fruit in IM arrest is not cumulative, and that fruit are only required to trigger arrest when the IM is competent to do so. Furthermore, our data show that it is only the fruits proximal to the IM at the time of arrest that are required to drive arrest, and we find no evidence that older fruit play a significant role in the arrest process. Collectively, our data can be used to refine the model to suggest that 6-8 fertile fruit proximal to the IM are the minimum required to trigger arrest in competent IMs, since 6-8 fruits per inflorescence are produced before arrest in both continuously de-flowered (**Fig 1B**) and in ‘late’ de-flowered plants (**Fig 3C**), once the treatment is ended.

**Figure 3.**
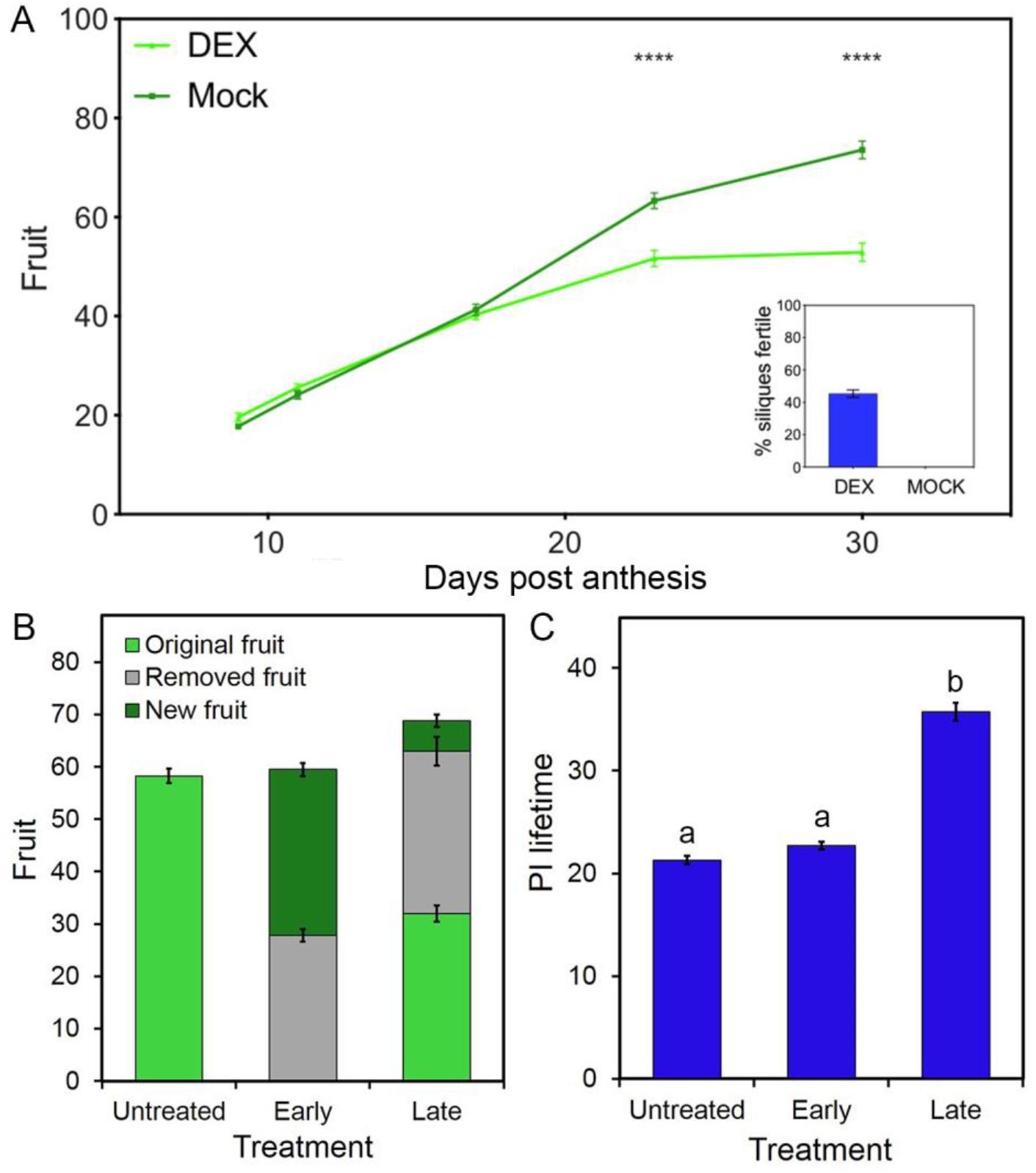
Proximal fruit drive arrest in inflorescence meristems. (**A**) Floral arrest is delayed by male sterility. Mock treated *MS1:MS-GR ms1-1* plants are fully sterile and do not undergo timely primary inflorescence arrest, behaving the same as *ms1-1* plants. However if fertility is restored by 25μm DEX treatment applied to fruit at 11 and 12 days post anthesis (dpa) of the first flower on the primary inflorescence, timely inflorescence arrest occurs. Application of DEX to fruit resulted in approximately 45% fertility, while mock-treated plants exhibited no fertility (inset). n= 9-12, bars indicate s.e.m. Stars indicate significance as determined by Sidak’s multiple comparisons following fitting of a mixed-effects model (**** = p <0.0001). (**B, C**) Effect of partial and differential fruit removal on inflorescence meristem arrest. In ‘early’ plants, open flowers were removed from the whole plant every 1-2 days until approximately 30 flowers had been produced on the primary inflorescence, following which they were allowed to flower normally. ‘Late’ plants were allowed to flower as normal until around 30 flowers had opened on the primary inflorescence; all subsequently-produced flowers were removed from the whole plant daily. (**B**) Shows the number of nodes produced by the PI in these treatments, coloured according to whether the flower was removed or fruit were produced before or after treatment. (**C**) Shows the inflorescence lifetime of the PI for these different treatments. *n* = 11-12, bars indicate s.e.m. Bars with the same letter are not statistically different from each other (ANOVA, Tukey HSD test).

We next questioned how fertile fruit signal to IMs. Previous work tentatively proposed that fruit communicate with IMs by a phytohormonal signal, although provided no clear evidence supporting this [1,2,3]. Given the previously described role of auxin export in co-ordinating the activity of IM in Arabidopsis [6,7], and the high levels of auxin known to be produced in fruits and seeds in many species [8,9,10,11], we tested whether auxin itself might be the mobile ‘fruit-to-meristem’ (F-M) signal. We firstly tested whether exogenous application of auxin to fruits of an infertile mutant could re-initiate the timely arrest of the PI. We treated sterile fruit in the *ams* mutant, which like *ms1* fails to go through normal floral arrest [12], with the auxin analog NAA from 6 dpa. This resulted in earlier IM arrest with the PI producing ~50 fruit, compared to ~80 in mock-treated plants (**Fig. 4A**). In auxin-treated *ams* plants, IM arrest phenocopies the normal ‘bud cluster’ morphology associated with the arrest of wild-type IMs (**Fig. 4C**). As expected, auxin treatment early in flowering does not induce IM arrest, and is only able to do so when IMs become competent to arrest (**Fig. 4A**). When we applied NAA to the uppermost 10 sterile fruit of *ams* individuals at 20 dpa (and to any fruit subsequently formed in the following 3 days), this rapidly induced a normal IM arrest (**Fig. 4B**) through the treatment of relatively few (~18) fruit (**Fig. 4B**), consistent with the role of proximal fruit in triggering IM arrest.

**Figure 4.**
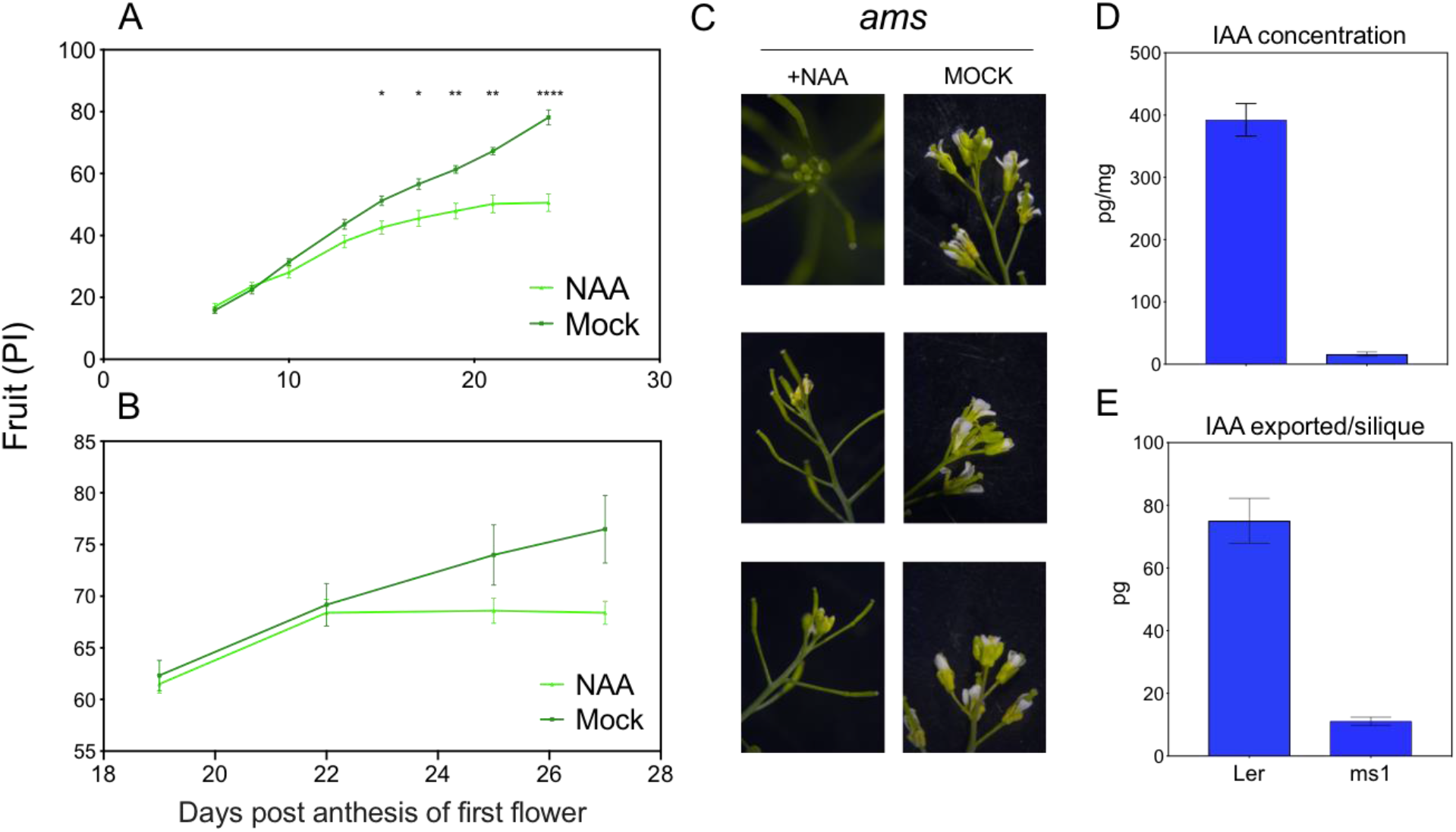
Auxin export from fruit triggers inflorescence meristem arrest. A) Number of fruit produced on the PI of male-sterile *ams* plants upon application of either 5mg/g NAA in lanolin or a mock treatment consisting of lanolin and DMSO. Fruit counts and lanolin treatment were performed every 2-3 days, starting from 6 days post anthesis (dpa) of the first flower on the primary inflorescence. n = 7-12, bars indicate s.e.m. B) Number of fruit produced on the PI of male-sterile *ams* upon application of 5mg/g NAA in lanolin or mock as in (A). Fruit counts and lanolin treatment were performed every day, starting from 20 days post anthesis dpa. n = 6-10, bars indicate s.e.m. Asterisks in both A and B indicate significance as determined by Sidak’s multiple comparisons following fitting of a mixed-effects model; * = <0.05; ** = <0.01; *** = 0.001; **** = 0.0001. C) Apical arrest phenotypes of NAA and mock treated *ams* plants. D) Quantification of auxin content in 6 dpa fertile (L*er*) and sterile (*ms1*) Arabidopsis fruits. n=5, bars indicate SD. E) Quantification of auxin eluted from fertile and sterile Arabidopsis fruits. n= 5, bars indicate SD.

As the previous data showed that auxin was sufficient to induce timely arrest of IM in infertile fruits, we next explored whether fertile fruits could provide a source for auxin. Previous work in Arabidopsis has identified a curve of hormone production in developing fruit, with a peak in auxin content at 6 dpa [11]. To assess whether fertilisation increases the auxin content of Arabidopsis fruit, sterile (*ms1-1*) and fertile (L*er*) fruit were sampled at 6 dpa, and auxin levels were quantified using LC-MS/MS. This analysis showed that auxin levels are greatly increased when fruit are fertile. The auxin content (pmol/g of tissue) of fertile fruit at this timepoint was 392pg/mg, while sterile fruit contained 16pg/mg of auxin (**Fig. 4D**). Fertile fruit therefore represent a strong potential auxin source, and the difference relative to sterile fruit is further amplified by an approximate 10-fold weight difference between them (**Fig. S2**). We next ascertained whether fertile fruit indeed transport auxin into the stem, enabling auxin to act as a F-M signal, by collecting auxin exported from the pedicels of 6 dpa fertile fruit from the PI. We found that individual fertile fruit export ~75pg of auxin in 21 hours, which is 7.5 fold higher than equivalent sterile fruit (**Fig. 4E**). Given that the equivalent pool of mobile auxin collected from the associated inflorescence stem is ~100-200pg [7], it is clear that individual fertile fruit make a very significant contribution to auxin levels in the inflorescence stem, supporting a model in which auxin export provides the F-M signal.

Our research provides clearer understanding of the process of floral arrest in Arabidopsis, and the regulatory mechanisms that govern it. We show that floral arrest arises from the uncoordinated local arrest of IMs, rather than a globally coordinated arrest, and that quasi-synchronicity of floral arrest is a natural consequence of the quasi-synchronous IM activation. We show that IMs will only arrest when they become temporally-competent to do so, which is likely a reflection of the developmental age of the IM. Our work thus complements the recent work of Balanzà et al [3] who showed that age-related up-and down-regulation of the FRUITFUL and APETALA2 transcription factors was associated with delayed floral arrest. FRUITFUL and APETALA2 are thus likely to be key factors determining the competence of IMs to arrest.

We have shown that auxin exported from fruits may act as the F-M signal. Wuest et al [2] showed that arrested IMs have a similar transcriptome to dormant IMs, suggesting IM activity is a reversible state, and that IM arrest might occur by the inverse mechanism to IM activation. Based on the canalisation model of bud activation [6,7,13], we therefore propose that competent IMs are tipped back into a dormant state because they become out-competed for canalisation of auxin export by the auxin exported from proximal fruit. This in turn suggests that the arrest-competent state is associated with a rapid loss of auxin source strength in the IM (Fig. 5). Our work thus potentially expands the canalization framework to a new developmental process, but more work will be needed to test and model these ideas. Overall, our model refines Hensel et als’ GPA model [1], and provides a mechanistic framework which would allow for the duration of flowering to be tailored to match local climatic conditions, whilst also containing a key checkpoint so that flowering only ceases if fertile fruit have recently been made. This paves the way to provide understanding of the end-of-flowering syndromes in other species, which in turn has potential impact for extending and maximising future crop yields.

**Figure 5.**
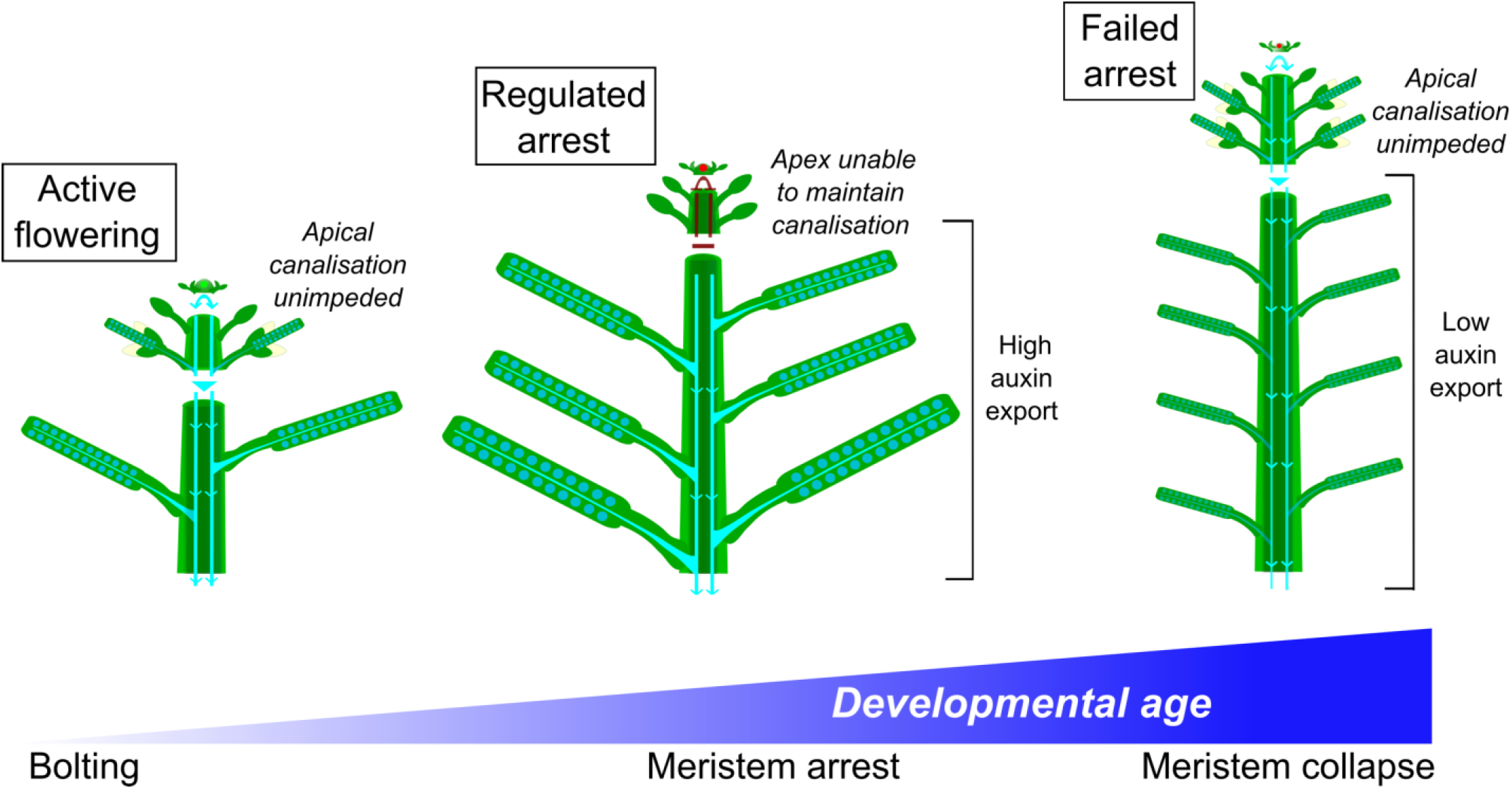
Model for induction of floral arrest. Initially, IMs are young (green) and can canalize to the polar auxin transport stream (PATS, blue). After a temporally-defined period of flowering, IMs reach a critical age and become capable of arrest (red). In the presence of ca. 6-8 proximal fertile fruit, which actively export auxin into the PATs, the meristem is unable to canalize. This induces floral arrest, similar to bud dormancy. If fruit are sterile (or removed), the auxin export capacity from proximal fruit is significantly reduced. This allows flowering to be prolonged beyond the point of the IM becoming arrest-competent, as it can still canalize to the PATS. Fertilisation, or auxin application, at this point rapidly induces arrest. If no fertilisation occurs, the meristem ultimately collapses.

## MATERIALS & METHODS

### Plant growth conditions

Plants for phenotypic and microsurgical experiments were grown on John Innes compost, under a standard 16 h/8 h light/dark cycle (20°C/16°C) in controlled environment rooms with light provided by white fluorescent tubes at a light intensity of ~120μmol/m^2^s^−1^. Plants for hormone profiling, dexamethasone application and hormone application experiments were grown on John Innes No.3 compost under the same light/dark cycle but at 22°C/18°C, with light provided by fluorescent tubes at an intensity of ~150μmol/m^2^s^−1^.

### Plant materials

Arabidopsis wild-types Col-0 and L*er* were used as indicated. The following lines have previously been described before; *ms1-1* (L*er* background) [14]; *AMS:AMS-GR ams* (Col-0 background, *ams* is SALK_152147) [12]; *MS1:MS1-GR ms1-1* (L*er* background) [15].

### Phenotypic assessments

We used the following nomenclature. The primary embryonic shoot apex gives rise to primary leaves and eventually forms the primary inflorescence. Flowering branches that form from axillary buds in the axils of primary leaves are secondary inflorescences. Secondary inflorescences formed from primary cauline leaves are cauline inflorescences (e.g. C1, C2, etc.), those from primary rosette leaves are rosette inflorescences (R1, R2, etc.). Secondary inflorescences are numbered in rootward fashion, such that C1 is the uppermost cauline inflorescence, and R1 the uppermost rosette inflorescence. Branches that form from secondary inflorescences are tertiary inflorescences, etc, and are named after the parental branching system in rootward fashion (e.g. C2.1 = uppermost tertiary branch on the second cauline inflorescence).

For the timing data in Fig. 1D, 2A, 2B, 2C and 3D, plants were assessed daily until visible flower buds were present at the shoot meristem. This date of floral transition was recorded, and plants were assessed daily as appropriate for IM activation (scored when buds were longer than 1mm) and IM arrest (scored when there were no more open flowers on the IM). For fruit counts in Fig. 2D, 2E, 3C, the number of inflorescences was counted, and the number of fruits on each inflorescence recorded (or the number of fruits removed). Fruit counts were made at final arrest unless otherwise stated.

For the DEX-induction experiment, *MS1:MS1-GR ms1-1* plants were treated with either a solution consisting of 10ml distilled water, 25μM Dexamethasone (from a 25mM stock in ethanol), and 2μl Silwet-77, or a mock containing the same but with only ethanol. Treatments were carried out at 11 and 12 dpa and fruit number was subsequently counted at the time points indicated on the graph.

Following the arrest of the DEX-treated plants, the percent of fertility in all plants was evaluated counting the number of fruit which had extended.

### Micro-surgical experiments

Flower removal in Fig 1A-D, 3C-D was performed every 1 to 2 days by removing all open flowers on the plant between the stated time points.

### IAA metabolite quantification

For quantification of IAA and IAA metabolites, 6 dpa fruits were sampled from mature flowering (ca. 15-18 dpa) *ms1-1* and L*er* plants. Fruit age had been tracked by marking their corresponding flowers with thread at 6 days previously, at anthesis. For the export assay the same strategy was used, but following excision fruits were placed pedicel-down in closed PCR tubes containing 50ul 2.5mM sodium diethyldithiocarbamate buffer and incubated for 21h in a growth room. The samples were snap frozen in liquid nitrogen and stored at −80°C until analysis, either by GC-MS/MS as described in Prusinkiewicz et al 2009 (eluates) or by UHPLC-MS/MS as described in [16], where prior the UHPLC-MS/MS analysis the fruit tissues were extracted and purified according to [17].

### Hormone applications

For the 5mg/g NAA lanolin treatments, 50ul of either 100mg/ml stock solution in DMSO or just DMSO for the mock with 1ul of dye was added to 1g of molten lanolin (heated to 60°C) and subsequently shaken until completely incorporated. Enough of the paste to create a thin layer was then applied using a micropipette tip to the fruit. For the early/continual NAA application experiments, the application regimen began at 6 dpa of the first flower. For the late NAA application experiment, treatment was initiated at 20 dpa and only the top (i.e. proximal to the IM) 10 fruits, and any produced above these in the subsequent 3 days were treated. Treatments were conducted at the same time as fruit number counts, indicated by the time points on the graphs.

### Experimental design and statistics

Samples size for each experiment are described in the figure legends. For plant growth experiments, each sample was a distinct plant. For auxin measurements, each sample was set of tissue pooled from multiple plants; each sample was distinct. For data analysis, normality was not assumed, and a general linear model was used to determine the most appropriate statistical test.

## ACKNOWLEDGEMENTS

AW is supported by BBSRC DTP grant BB/M008770/1. KL and JS are supported by the Knut and Alice Wallenberg Foundation (KAW), the Swedish Governmental Agency for Innovation Systems (VINNOVA) and the Swedish Research Council (VR). We also thank Roger Granbom for technical assistance and the Swedish Metabolomics Centre (http://www.swedishmetabolomicscentre.se/) for access to instrumentation.

## AUTHOR CONTRIBUTIONS

CW, AW, JS, KL performed experiments and analysed the data. TB, AB & ZW designed the study. All authors contributed to writing the manuscript.

---

**Supplementary Table 1.** Details of experiments used for flowering duration and fruit assessments. Means are presented for the untreated controls in each experiment ± standard deviation. All experiments were performed under similar long day conditions (16h day/8h night), grown in either a greenhouse with supplementary lighting (GH) or a walk-in controlled environment chamber (WI).

| Experiment number | GH or WI | Floral duration | PI duration | Initial fruit count | RF fruit count |
| --- | --- | --- | --- | --- | --- |
| 1 | WI | 26.4 ± 2.0 |  |  |  |
| 2 | WI | 28.1 ± 2.5 | 21.3 ± 1.3 | 515 ± 128 |  |
| 3 | WI | 25.4 ± 2.6 | 19.3 ± 1.3 |  |  |
| 4 | WI | 24.4 ± 0.9 | 19.4 ± 1.3 |  |  |
| 6 | WI | 27.1 ± 2.9 | 22.3 ± 1.1 | 379 ± 62 | 513 ± 75 |
| 7 | WI | 25.4 ± 1.4 | 20.1 ± 1.0 |  |  |
| 8 | WI |  |  | 184 ± 30 | 208 ± 27 |
| 9 | GH |  |  | 307 ± 69 |  |
| 10 | GH |  |  | 648 ± 66 |  |
| 11 | GH |  |  | 345 ± 119 |  |
| 12 | WI |  |  | 398 ± 81 |  |
| 13 | WI |  |  | 454 ± 81 |  |
| 14 | WI |  |  | 330 ± 108 |  |
| 15 | WI |  |  | 501 ± 101 | 571 ± 149 |
| 16 | WI |  |  | 352 ± 129 | 483 ± 173 |
| 17 | WI |  |  | 241 ± 65 | 301 ± 86 |

**Supplementary Figure 1.**
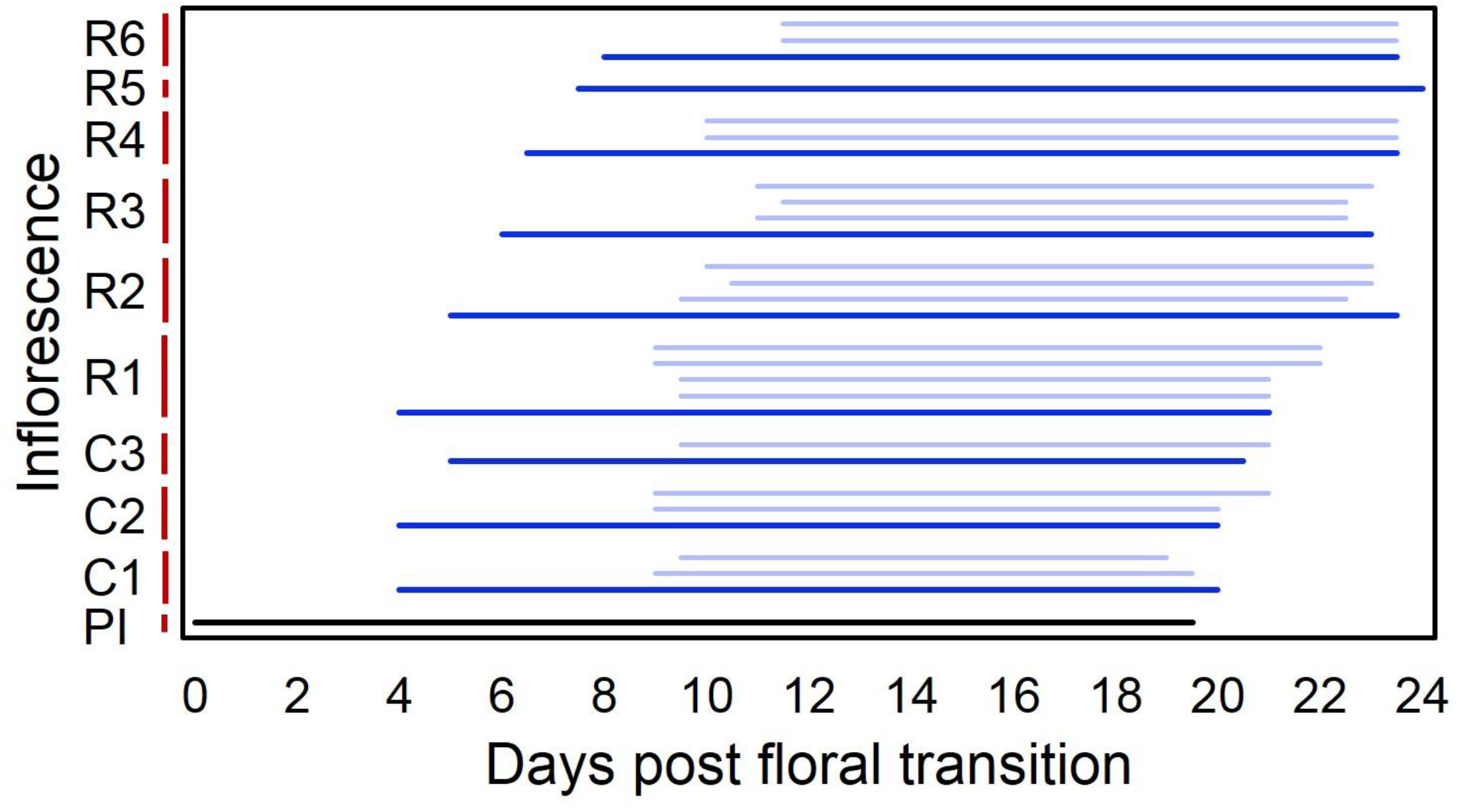
Complete dataset for data shown in Figure 2A, 2B. Lifetime of individual inflorescences in Col-0, from inflorescence activation to arrest, shown relative to the time since bolting started. The primary inflorescence (‘PI’) is indicated in black, secondary cauline (C1, C2, etc.) and rosette (R1, R2, etc.) inflorescences in dark blue. Tertiary inflorescences are shown in light blue above their parent secondary inflorescence. Values are derived from analysis of 8 plants. Each inflorescence lifetime is the mean of 3-8 plants, depending on which plants had which inflorescence type. Any inflorescence type occurring on two or fewer plants was excluded from analysis.

**Supplementary Figure 2.**
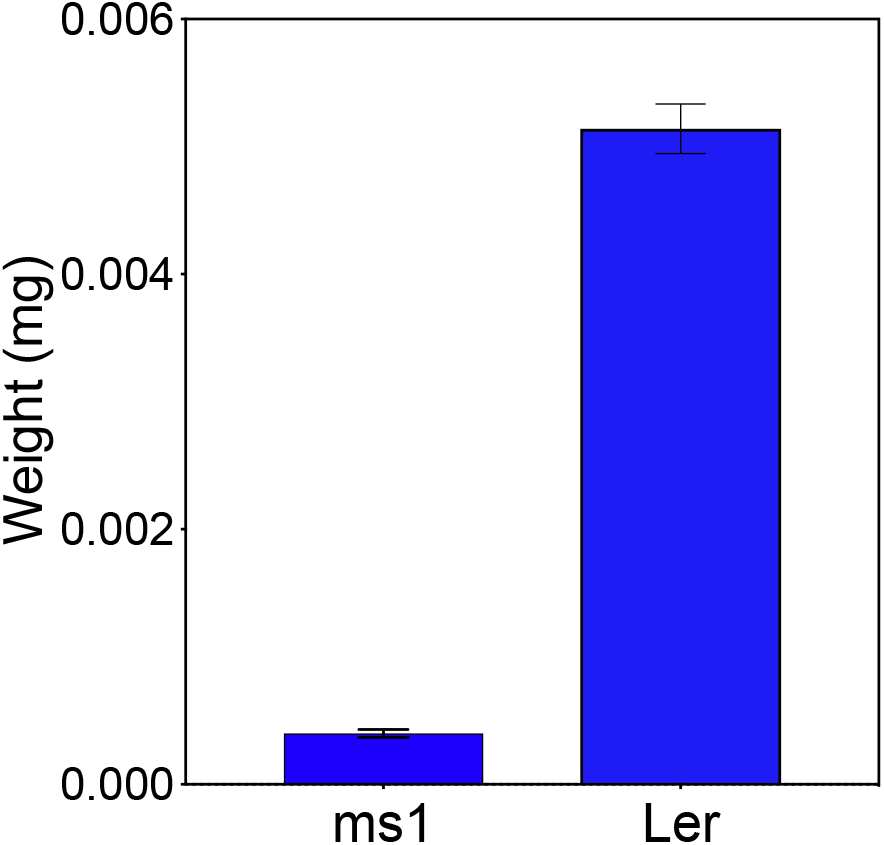
Weight of fertile versus sterile Ler and *ms1-1* siliques, as measured at days post anthesis (dpa) of the first flower on the primary inflorescence. N = 5, error bars indicate s.e.m.

